# Precise levels of Nectin-3 and an interaction with Afadin are required for proper synapse formation in postnatal visual cortex

**DOI:** 10.1101/870691

**Authors:** Johanna Tomorsky, Philip R. L. Parker, Chris Q. Doe, Cristopher M. Niell

## Abstract

**Background:** Developing cortical neurons express a tightly choreographed sequence of cytoskeletal and transmembrane proteins to form and strengthen specific synaptic connections during circuit formation. Nectin-3 is a cell-adhesion molecule with previously described roles in synapse formation and maintenance. This protein and its binding partner, Nectin-1, are selectively expressed in upper-layer neurons of mouse visual cortex, but their role in the development of cortical circuits is unknown.

**Methods:** Here we block Nectin-3 expression (via shRNA) or overexpress Nectin-3 in developing layer 2/3 visual cortical neurons using *in utero* electroporation. We then assay dendritic spine densities at three developmental time points: eye opening (postnatal day (P)14), one week following eye opening after a period of heightened synaptogenesis (P21), and at the close of the critical period for ocular dominance plasticity (P35).

**Results:** Knockdown of Nectin-3 beginning at E15.5 or ∼P19 increased dendritic spine densities at P21 or P35, respectively. Conversely, overexpressing full length Nectin-3 at E15.5 led to decreased dendritic spine densities when all ages were considered together. Interestingly, an even greater decrease in dendritic spine densities, particularly at P21, was observed when we overexpressed Nectin-3 lacking its Afadin binding domain, indicating Afadin may facilitate spine morphogenesis after eye opening.

**Conclusion:** These data collectively suggest that the proper levels of Nectin-3, as well as the interaction of Nectin-3 with Afadin, facilitate normal synapse formation after eye opening in layer 2/3 visual cortical neurons.

## Background

The development of mature brain circuits requires the selective formation, elimination, and strengthening of synaptic connections between neurons [1,2]. This process requires the timed expression and spatial localization of proteins that are just beginning to be elucidated [3–5]. Many studies indicate that cell-adhesion molecules are involved in the formation and strengthening of synapses, though much is unknown about their specific roles in the maturation of various brain circuits [6–11]. The number and strength of synapses found on visual cortical neurons, inferred from dendritic spine density and shape, depend on visual experience and developmental stage [1,12–19]. During the postnatal development of visual cortex, neurons experience an early increase in dendritic spine densities following eye opening [18,20,21]. This period of heightened synaptogenesis is followed by the critical period for ocular dominance plasticity (ODP, postnatal day (P)21-P35), which is associated with experience-dependent synaptic refinement and pruning [2,4,19,20,22]. Dendritic spine remodeling provides a mechanism for synapses to alter their strength and number based on experience, e.g. through activity-dependent synaptic plasticity, and is a critical component of normal cortical development [12,14–16,23,24]. Moreover, each cortical layer has a distinct timeline for the development of its unique visual response properties [25], likely a result of layer-specific synapse remodeling. Here, we sought to characterize the role of the cell-adhesion molecule Nectin-3 in spine density profiles in developing layer 2/3 (L2/3) visual cortical neurons.

The nectins are a group of immunoglobulin superfamily cell-adhesion molecules found to stabilize synapses at specialized structures called puncta adherentia junctions (PAJs) [26–29]. Trans-synaptic nectins and cadherins, together with their intracellular binding partners (Afadin and catenins, respectively), help to form and remodel synapses over development (Figure 1a) [11,30]. For example, axonal (presynaptic) Nectin-1 in dentate granule cells specifically binds to dendritic (postsynaptic) Nectin-3 in CA3 principal neurons at the stratum lucidum in hippocampus to help guide and stabilize developing synaptic connections between these neurons (Figure 1a) [11,31]. Other studies have implicated nectins in a variety of biological and disease states, including long-term memory formation, stress, taopathy, and mental retardation, indicating that nectins may have different functions in the development, aging, and maintenance of various brain circuits [32–36]. While previous work suggests that Nectin-1 and Nectin-3 may be involved in both the formation and maturation (stabilization) of synapses in hippocampus, their function in postnatal cortical development is unknown [11].

Here we identify a role for Nectin-3 in the development of visual cortical neurons by examining dendritic spine densities after manipulating neuronal Nectin-3 expression *in vivo*. Mammalian cortex has a conserved laminar structure, and Nectin-1 and Nectin-3 both have distinct upper layer expression patterns in postnatal mouse cortex [5,32,37]. This expression pattern suggests a potential role for Nectin-3 in the development of L2/3-specific visual response properties after eye opening [25]. We assessed the dendritic spine densities of L2/3 visual cortical neurons at three developmental time points (P14, P21, and P35) after *in utero* electroporation of plasmid constructs expressing Nectin-3 shRNA, full-length Nectin-3, or Nectin-3 lacking the four amino acid Afadin-binding domain (Nec3^Δafadin^) [38–40]. Nectin-3 knockdown by shRNA beginning at embryonic day (E)15.5 or, using CaMKII-Cre transgenic mice [41], ∼P19 increased dendritic spine densities at P21 or P35, respectively. In contrast, overexpressing full-length Nectin-3 produced an overall decrease in dendritic spine densities. Interestingly, overexpressing Nec3^Δafadin^ produced an even greater decrease in dendritic spine densities than full-length Nectin-3, particularly at P21. Our results indicate that the neuronal regulation of synaptic Nectin-3, as well as an interaction with Afadin, may mediate both the formation and selective removal of synapses on L2/3 visual cortical neurons during critical periods of development.

**Figure 1.**
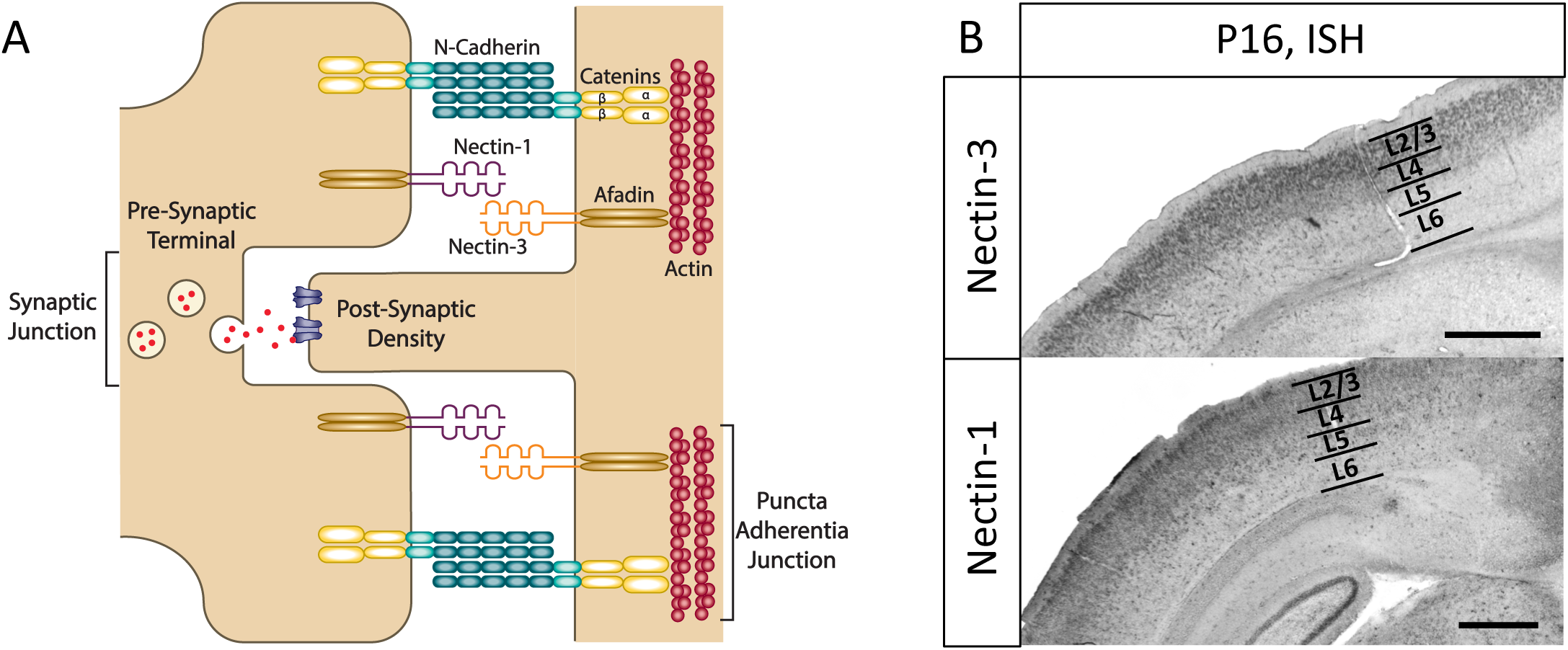
Model of synaptic Nectin-3 and Nectin-1 interactions and expression in upper layers of visual cortex. **(a)** Nectin-1 and Nectin-3 have been shown to bind at puncta adherentia junctions (PAJs) in hippocampus and interact with actin secondarily through their binding partner Afadin [11,28,31]. Nectin and Afadin have also been shown to associate with the N-cadherin/catenin complex at PAJs [11,27–29,31,39]. Synaptic PAJs are stabilizing sites of adhesion between axons and dendrites and are distinct from synaptic junctions, which are the sites of neurotransmission [11,28]. **(b)** *In situ* hybridizations (ISH) to Nectin-3 and Nectin-1 demonstrate enriched expression in upper cortical layer neurons in V1 at P16 (5x objective, scale bar = 500 µm).

## Methods

### *In situ* hybridization

The expression patterns of Nectin-1 and Nectin-3 were assayed by nonradioactive colorimetric RNA *in situ* hybridization, using solutions and probes as previously described [5,42–44]. Briefly, animals were perfused and brains were fixed overnight in 4% PFA and then cryoprotected in a 30% sucrose solution. Brains were then cryosectioned to a thickness of 30 µm, mounted on Superfrost Plus slides (Fisherbrand), and stored at -80 °C until use. 30 µm sections were brought to room temperature, washed in PBS and acetylated [44]. Slides were then pre-hybridized in hybridization solution in a humidity chamber for 2 h at 70 °C. The riboprobes to *Pvrl1* and *Pvrl3* (the genes encoding Nectin-1 and Nectin-3 proteins) were generated with dig-labeled nucleotides and SP6 RNA polymerase using probe sequences and protocols described by the Allen Brain Institute [37,44,45]. Riboprobes were diluted in hybridization solution to a concentration of 1–2 ng/µL, and slides were hybridized overnight at 70 °C with each probe. Slides were then washed (at 70 °C) and blocked for 1 h (at room temperature) before incubating overnight at 4 °C in anti-dig sheep Fab fragments conjugated to alkaline phosphatase (AP; Roche No. 11093274910) diluted 1:2500 in blocking solution. Slices were then washed at room temperature with MABT buffer and then AP staining buffer, after which 3.5 µL/mL NBT, 2.6 µL/mL BCIP, and 80 µL/mL levamisole in AP staining buffer was applied. The AP colorimetric reaction was observed closely as it developed for 3–48 h at 37 °C and was stopped by washing twice with PBS (0.1% Tween-20) and twice with deionized H2O. Slides were then dehydrated in graded ethanols and mounted using Permount Mounting Medium (Fisher).

### Design of Cre-dependent shRNA plasmids for *in utero* electroporation

Nectin-3 shRNA constructs were designed using previously published siRNA sequences [40]. These 19 bp sequences were used to design shRNA sequences for cloning into a pSico vector [40,46]. The Nectin-3 and scramble shRNA oligos used for cloning were as follows:

Pvrl3 shRNA1 sense oligo:

TGGCCGGATTCTTTAATTGATTCAAGAGTCAATTAAAGAATCCGGCCTTTTTTC

Pvrl3 shRNA1 antisense oligo:

TCGAGAAAAAAGGCCGGATTCTTTAATTGACTCTTGAATCAATTAAAGAATCCGGCC A

Pvrl3 shRNA2 sense oligo:

TGTTTATTGGCGTCAGATAATTCAAGAGATTATCTGACGCCAATAAACTTTTTTC

Pvrl3 shRNA2 antisense oligo:

TCGAGAAAAAAGTTTATTGGCGTCAGATAATCTCTTGAATTATCTGACGCCAATAAACA

Scr sense oligo:

TGCTACACTATCGAGCAATTTTCAAGAGAAATTGCTCGATAGTGTAGCTTTTTTC

Scr antisense oligo:

TCGAGAAAAAAGCTACACTATCGAGCAATTTCTCTTGAAAATTGCTCGATAGTGTAGCA

Cloning into pSico was modified from methods previously described [46]. Briefly, restriction enzymes XhoI and HpaI were used to digest the pSico vector (Addgene plasmid #11578). Digested vector RNA was then dephosphorylated using shrimp alkaline phosphatase (Roche) for 60 min at 37 °C to prevent re-ligation of the vector, followed by deactivation at 65 °C for 15 min. 1 µL of 100 µM sense and antisense shRNA oligos (synthesized by IDT) were then annealed and phosphorylated using T4 PNK (1µL 10X T4 Ligation Buffer (NEB), 6.5 µL H2O, and 0.5 µL T4 PNK (NEB)). Annealing was performed in a thermocycler set to 37 °C for 30 min and then 95 °C for 5 min, followed by a ramp down to 25 °C at 5 °C/min. Vector and insert were then ligated with Quick Ligase (NEB) using manufacturers protocols. Ligated plasmid was then treated with Plasmid-Safe exonuclease (Lucigen), to prevent unwanted recombination products, and used to transform NEB Stable Competent *e. coli*. Positive colonies were grown in LB + amp., after which plasmid DNA was extracted (QIAprep Spin Miniprep Kit) and sequenced using the pSico sequencing primer: CAAACACAGTGCACACAACGC [46]. Plasmid DNA verified to contain the shRNA insert was then prepped for electroporation from 200–400 µL of cultured *e. coli* using a NucleoBond Midi or Maxi EF kit (Clontech) and eluted to a concentration of 5–10 µg/µL in TE.

### Expression constructs for *in utero* electroporation

A Nectin-3 overexpression construct was created by modifying a pCag-iCre expression vector (Addgene plasmid # 89573). Nectin-3 alpha was PCR amplified (KOD hot start DNA polymerase, Sigma-Aldrich) from a mouse brain cDNA library using the forward primer: GTTGAGGACACGCGCG and reverse primer: CTGTTAGACATACCACTCCCTCC. Amplified DNA was run on a gel and a band the length of Nectin-3 (∼1800 bp) was cut from the gel and purified (Qiaquick Gel Extraction Kit, Qiagen). Sequences were then amplified (KOD hot start DNA polymerase) using nested primers with and without a FLAG-tag and containing restriction sites for MluI and Not1. Primer sequences were as follows:

Nectin-3 F: TAAGCA-ACGCGT-GCCACCATGGCGCGGACCCCG

Nectin-3 R: TGCTTA-GCGGCCGC-TTA-GACATACCACTCCCTCCTG

Nectin-3 R Flag: TAAGCA-GCGGCCGC-TTA-CTTGTCGTCATCGTCTTTGTAGTC-GACATACCACTCCCTCCTG

Amplified DNA was then gel purified and digested using the above mentioned restriction enzymes. The pCAG-iCre plasmid was also digested with MluI and NotI to remove the iCre sequence from the vector. Digested vector and Nectin-3 were then ligated, used to transform *e. coli*, and prepped for electroporation (NucleoBond Midi or Maxi EF kit, Clontech). Positive clones were sequenced using pCag F: GCAACGTTGCTGGTTATTGT, and Bglob-pA R: TTTTTGGCAGAGGGAAAAGAT sequencing primers.

A previously published construct expressing a truncated Nectin-3 (lacking its 4 terminal amino acids) was kindly gifted by Dr. Cristina Gil-Sanz and Dr. Ulrich Mueller [40]. FLEX-tdTomato constructs (plasmid #51509 and #51505) and the Cre-expression plasmid (plasmid #51904) were obtained from Addgene. All constructs were designed to express the Nectin-3 alpha splice variant (with and without its 4 C-terminal amino acids).

### Western blot analysis of overexpression and shRNA constructs

The newly designed Nectin-3 overexpression construct was transfected (Lipofectamine Reagent, Thermo Fisher Scientific) into HEK-293 cells using manufacturer’s protocols. A western blot was then run using 10 ug of protein extracted from transfected cells, as previously described [47]. When stained with an anti-Nectin-3 antibody (Abcam Cat# ab63931, RRID:AB_1142394) western blot analysis showed greatly increased expression compared to endogenous HEK-293 expression (Figure 5b). The western blot was also stained with an alpha-tubulin antibody (Sigma-Aldrich Cat# T9026, RRID:AB_477593) to ensure similar overall protein levels between conditions. To determine whether the newly designed shRNA constructs were effective, Nectin-3 overexpression and shRNA plasmids were co-transfected into HEK-293 cells. To establish the efficiency of co-transfection, lipofectamine was first used to co-transfect GFP and RFP expressing constructs into HEK-293 cells using manufacturers protocols. Using a 4:3 ratio of GFP to RFP constructs, nearly 100% co-transfection was observed (data not shown). pSico Nectin-3-shRNA and scramble-shRNA constructs were altered before transfection to artificially replicate Cre-excision using site-directed mutagenesis (Q5 site directed mutagenesis kit: New England Biolabs). The GFP-stop sequence and one loxP site were removed from pSico using the following primers: F: CGCATAACTTCGTATAGTATAAATTA, R: AATTACTTTACAGTTAGGGTGAG. The new Nectin-3 shRNA and scramble shRNA plasmids did not require Cre for expression and were each co-transfected into HEK-293 cells with the Nectin-3 overexpression construct at a 4:3 ratio. Again, 10 ug of extracted protein was loaded onto a gel for western blot, and the blot was stained using Nectin-3 and alpha-tubulin antibodies. After staining, nectin-3 expression/blot intensity was much higher than that observed for alpha-tubulin. For this reason, images of the Nectin-3 bands and alpha-tubulin bands were adjusted separately to bring the intensities of each into dynamic range (a single adjustment was made for all bands of a given protein). Band intensities were then quantified using image-j, and an expression index for each condition was obtained by normalizing Nectin-3 intensity to alpha-tubulin intensity (technical replicates, N = 2, Figure 2b).

**Figure 2:**
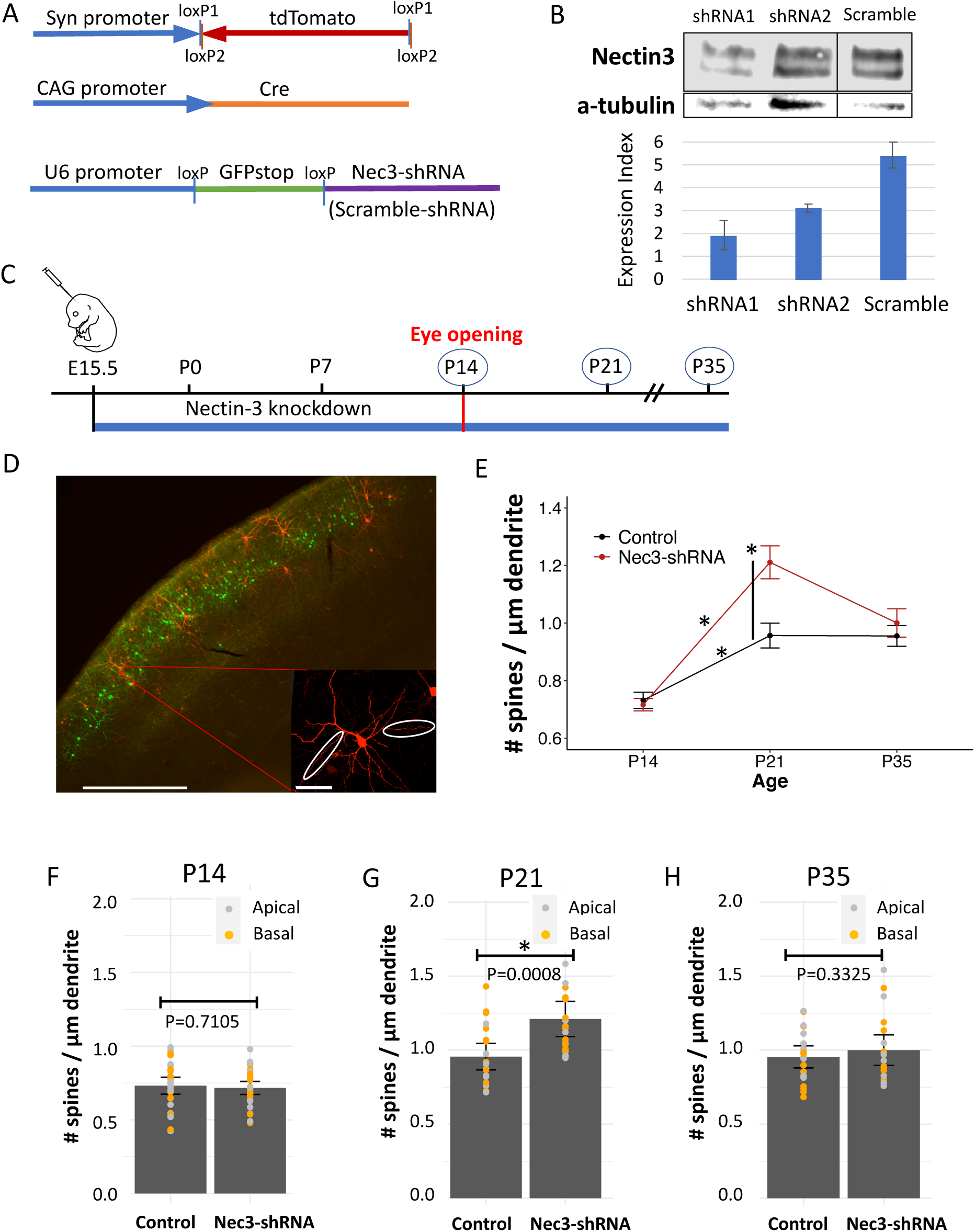
Nectin-3 knockdown at E15.5 increases spine density at P21. **(a)** Cre-dependent shRNA constructs to Nectin-3 (or scramble shRNA) contained loxP flanked GFP-stop sequences to prevent the expression of shRNA in the absence of Cre. shRNA constructs were co-electroporated with a Cre-expression plasmid and a Cre-dependent FLEX tdTomato construct. **(b)** Western blot showing Nectin-3 expression in HEK-293 cells after co-transfection of a Nectin-3 expression plasmid with either Nectin-3 or scramble shRNA. Nectin-3 expression/intensity normalized to α-tubulin is plotted (expression index, analyzed in ImageJ). **(c)** Expression of electroporated Nectin-3 or scramble shRNA occurred at E15.5. Dendritic spine densities were measured at three time points: P14, P21, and P35. **(d)** 10x image of neurons electroporated with a pSico shRNA construct at P21 in visual cortex. Cre-positive tdTomato-expressing cells assumed positive for shRNA expression can be observed surrounded by Cre-negative GFP-expressing cells (scale bar = 500 µm). Inset: Neurons were chosen in V1 (40x image, scale bar = 50 µm), and apical and basal dendrites, at least one branch away from the soma, were imaged at 63x for spine counting. **(e)** Differences in dendritic spine densities (pooled apical and basal dendrites) between neurons electroporated at E15.5 with a scrambled control shRNA or shRNA to Nectin-3. Error bars denote standard errors of the mean. From left to right, asterisks denote significant differences (95% confidence level after Bonferroni correction) between time points (control, P14-P21; *p* = 0.00027, and Nec-3 shRNA, P14-P21; *p* = 4.545e-09) or between conditions at the P21 time point (*p* = 0.0008, Additional File 1). **(f)** Nec3-shRNA and control dendritic spine densities at P14. Apical (grey) and basal (orange) dendritic spine densities are plotted along with error bars denoting the standard errors of the mean. **(g)** Nec3-shRNA and control dendritic spine densities at P21. Significance is denoted by ‘*’. **(h)** Nec3-shRNA and control dendritic spine densities at P35.

### *In utero* electroporation of plasmid DNA

All experimental protocols were approved by the University of Oregon Institutional Animal Care and Use Committees, in compliance with the National Institutes of Health guidelines for the care and use of experimental animals. *In utero* electroporation was performed at E15.5 to target L2/3 pyramidal neurons, as previously described [48,49]. For shRNA knockdown using pSico, a solution of 2 µg/µL pSico-shRNA plasmid, 1.5 µg/µL FLEX-tdTomato plasmid, and 0.02 µg/µL Cre-expression plasmid was prepared in PBS (pH 7.2 for injections). In addition, 0.1% Fast Green dye was used to visualize plasmid DNA as it entered the ventricle with injection. For overexpression experiments using either full-length or truncated Nectin-3, a solution of 0.5 µg/µL of expression vector, 1ug/µL FLEX-tdTomato plasmid, and 0.01 µg/µL Cre-expression plasmid was prepared in PBS and 0.1% Fast Green.

Timed pregnancies were set up overnight between a hybrid strain of WT female mice (F1 cross of C57BL/6J and 129S1/SvlmJ, Jax) and either the same strain of WT male mice or CaMKII-Cre homozygous transgenic male mice (Jax 005359). The day the plug was observed was designated embryonic day 0.5 (E0.5). 15.5-day pregnant mice were anesthetized with 2% isofluorane (0.8% O2) for the duration of the surgery. A small incision was made in the abdomen of pregnant females and the uterus was pulled out of the abdominal cavity. Approximately 1 µL of plasmid solution was injected through the uterus and scull into the lateral ventricle of E15.5 embryos [49]. Visual cortex was then targeted with an electrical pulse through tweezer-type electrodes using five, 45 V, 100 ms pulses at a 1 s interval. The uterus was then placed back in the abdominal cavity, the mouse was sutured, and allowed to recover. Animal health was monitored daily after surgery until pups were born (∼4 days later). Electroporated mice were perfused as previously described [50] at P14, P21, or P35, and brains were prepared for immunohistochemistry.

### Immunohistochemistry

tdTomato fluorescence in electroporated neurons was amplified by immunohistochemistry (IHC) before imaging. Tissue preparation for IHC was performed as previously described [50]. Briefly, brains were perfused and then fixed overnight in 4% PFA (1 x PBS), after which brains were immersed in 30% sucrose (1 x PBS) for 24–48 hours. Brains were then sliced on a vibratome to a thickness of 80 µm, placed in cryoprotectent solution (30% sucrose, 1% polyvinyl-pyrrolidone, 30% ethylene glycol in 0.1 M PB), and stored at -20 °C. IHC was performed on free floating sections as previously described [50]. Briefly, sections were washed 3 x 10 min in a 0.7% glycine solution (in PBS) and then blocked for 1–3 h in PBST (0.3% Triton-X in PBS) containing 5% goat and 5% donkey serum. Slices were then transferred to a primary antibody solution containing 1.5 µL/mL of rabbit anti-RFP (Rockland Cat# 600-401-379, RRID:AB_2209751) and incubated overnight at 4 °C. The next day, slices were washed 1 x 10 min in PBST and then 3 x 10 min in PBS. Next, slices were transferred to a solution of 4 µL/mL of anti-rabbit Alexa Fluor 555 (Thermo Fisher Scientific Cat# A-21429, RRID:AB_2535850) secondary antibody in PBST and incubated at room temperature for 3 h. Slices were then washed for 10 min in PBST at room temperature, followed by an overnight wash in PBS at 4 °C. The next day, slices were washed an additional 2 x 10 min in PBS, treated with DAPI (4′, 6-diamidino-2-phenylindole), and mounted with VECTASHIELD mounting media (Vector Labs).

### Microscopy and spine counting

*In situ* hybridizations were imaged using an EC Plan-NEOFLUAR 5x/0.16 objective on a Zeiss Axio Imager.A2 wide field epifluorescence microscope having an X-Cite 120Q LED excitation lamp and a Zeiss AxioCam MRm 1.4-megapixel camera. ZEN lite imaging software (2012) was used to view images, and Adobe Photoshop CS6 was used for background removal and color processing of images.

Coronal sections of electroporated brain tissue were checked for fluorescent neurons in V1 using the Zeiss Axio Imager.A2 microscope and an EC Plan-NEOFLUAR 2.5x/0.085 objective. Immunohistochemistry was performed on sections confirmed to have electroporated cells. The location of electroporated neurons was determined using DAPI staining and a mouse brain atlas [51]. Neurons located V1 L2/3 were imaged on a ZeissLSM700 confocal microscope using Zen software. Images of secondary (at least one branch away from the soma) apical and basal dendrites were analyzed. Secondary apical dendrites directly extending laterally from the primary apical dendrite were preferentially selected (as opposed to apical tufts). High resolution images of dendrites for spine counting were taken using a Plan-Apochromat 63x/1.40 Oil DIC objective with 1.1–1.3x zoom, a speed of 8, averaging of 2, z-resolution of 0.3 µm, and variable laser intensities to capture dendritic spines. Spines in high resolution images of dendrites were counted manually using the open source FIJI image analysis software and the multipoint tool. Neurite lengths were measured using the ‘simple neurite tracer’ plugin.

### Statistical analysis

In this study, apical and basal dendrites from a single cell were often counted and used as individual measurements of dendritic spine density. At least 9 cells per group, or 18 dendrites, were analyzed to estimate average dendritic spine densities. The metadata for all experimental groups is included in Additional File 1 (Metadata sheet), including *p* values from individual t-tests examining differences between apical and basal dendrites. To account for possible cell-specific effects, we used a linear mixed model to analyze our data where ‘cell’ was included as a random effect. An ANOVA was then performed on modeled data using the Kenward-Rodger method to determine degrees of freedom. Analyses were performed in R using the ‘lme4’ package. Individual comparisons between each experimental condition (Nec3-shRNA, Nec3-OE, Nec3^Δafadin^, and control) were made at each age (P14, P21, and P35) using the following linear mixed model in R:

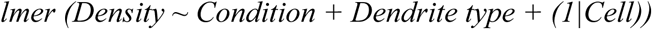

In this model, ‘condition’ and ‘dendrite type’ (apical or basal) are fixed effects, and ‘cell’ is included as a random effect. In addition, changes in dendritic spine densities between ages (P14 vs. P21 and P21 vs. P35) were analyzed for each condition using the following linear mixed model in R:

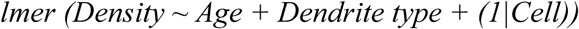

Unlike the first model, here ‘age’ is included as the first fixed effect. The data used in each comparison was first tested in R for normality and homoscedasticity using the Shapiro-Wilk normality test and Levene’s test for homogeneity of variance, respectively. If either test failed (p < 0.05), data was log2 transformed, after which the data for all but two comparisons passed both tests for normality and homoscedasticity (Additional File 1). Two comparisons with heteroskedastic data not corrected by log2 transformation had variances that differed by less than a factor of 2.5 and did not yield significant differences between conditions (Additional File 1, Additional File 2: Figure S1). All comparisons between conditions where animals were co-electroporated at E15.5 with a Cre plasmid (early manipulation of Nectin-3 expression) were considered a family, and a Bonferroni adjusted *p* value of 0.00192 was used to establish significant differences between groups (correction for a family of 26 comparisons). The experiment using a transgenic CamKII-Cre mouse for postnatal expression of Cre was considered an independent comparison, and a *p* value of 0.05 was used as a cut-off for significance (Figure 3).

**Figure 3.**
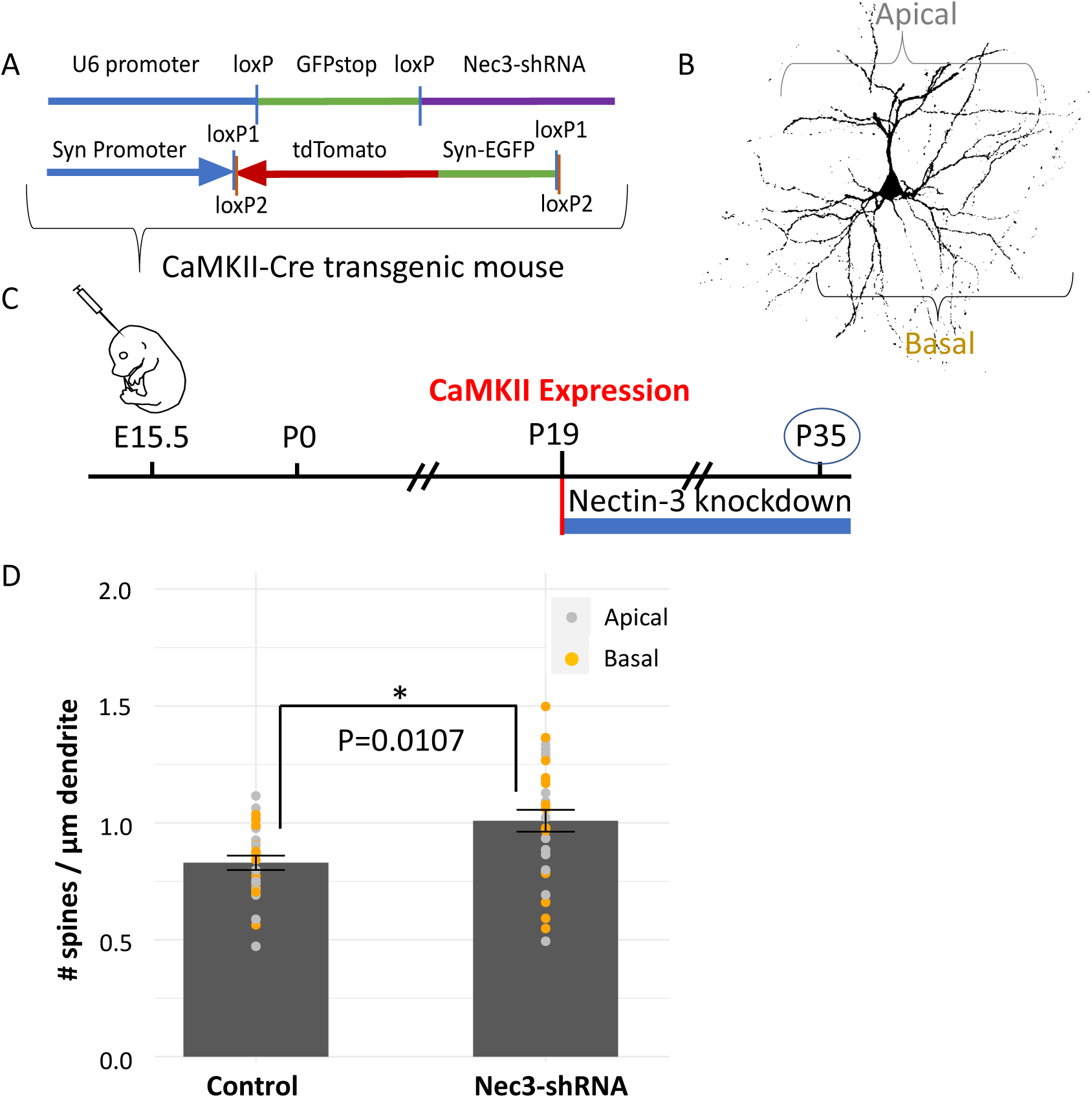
Nectin-3 knockdown at ∼P19 increases dendritic spine densities at P35. **(a)** A Cre-dependent shRNA construct to Nectin-3 (or scramble shRNA construct) was co-electroporated with a Cre-dependent FLEX tdTomato plasmid (also expressing synaptophysin-EGFP) into developing CaMKII-Cre transgenic mice. **(b)** Both apical and basal dendrites on CaMKII-Cre/shRNA/tdTomato positive neurons were imaged at P35 (40x image, scale bar = 50 µm). **(c)** Mice were electroporated at E15.5 to target developing L2/3 neurons, but Cre recombination is not observed until ∼P19 in the CaMKII-Cre mouse line used [41]. For this experiment, neurons develop normally until Nectin-3 knockdown at ∼P19. Mice were sacrificed and neurons were imaged at P35. **(d)** Dendritic spine densities are significantly higher at P35 when Nectin-3 is knocked down at ∼P19 using a CaMKII-Cre mouse. Apical and basal dendritic spine densities are indicated by grey and orange dots, respectively. Significance is denoted by ‘*’ (*p* = 0.0107, Additional File 1).

We performed three additional analyses to examine overall differences in dendritic spine densities between control (scramble shRNA) and either Nec3-shRNA (Figure 2), Nec3-OE (Figure 4), or Nec3^Δafadin^ (Figure 5) conditions. Data collected at all ages was included in these analyses. For these comparisons, both ‘condition’ and ‘age’ were included as fixed effects in our model with a potential interaction:

**Figure 4:**
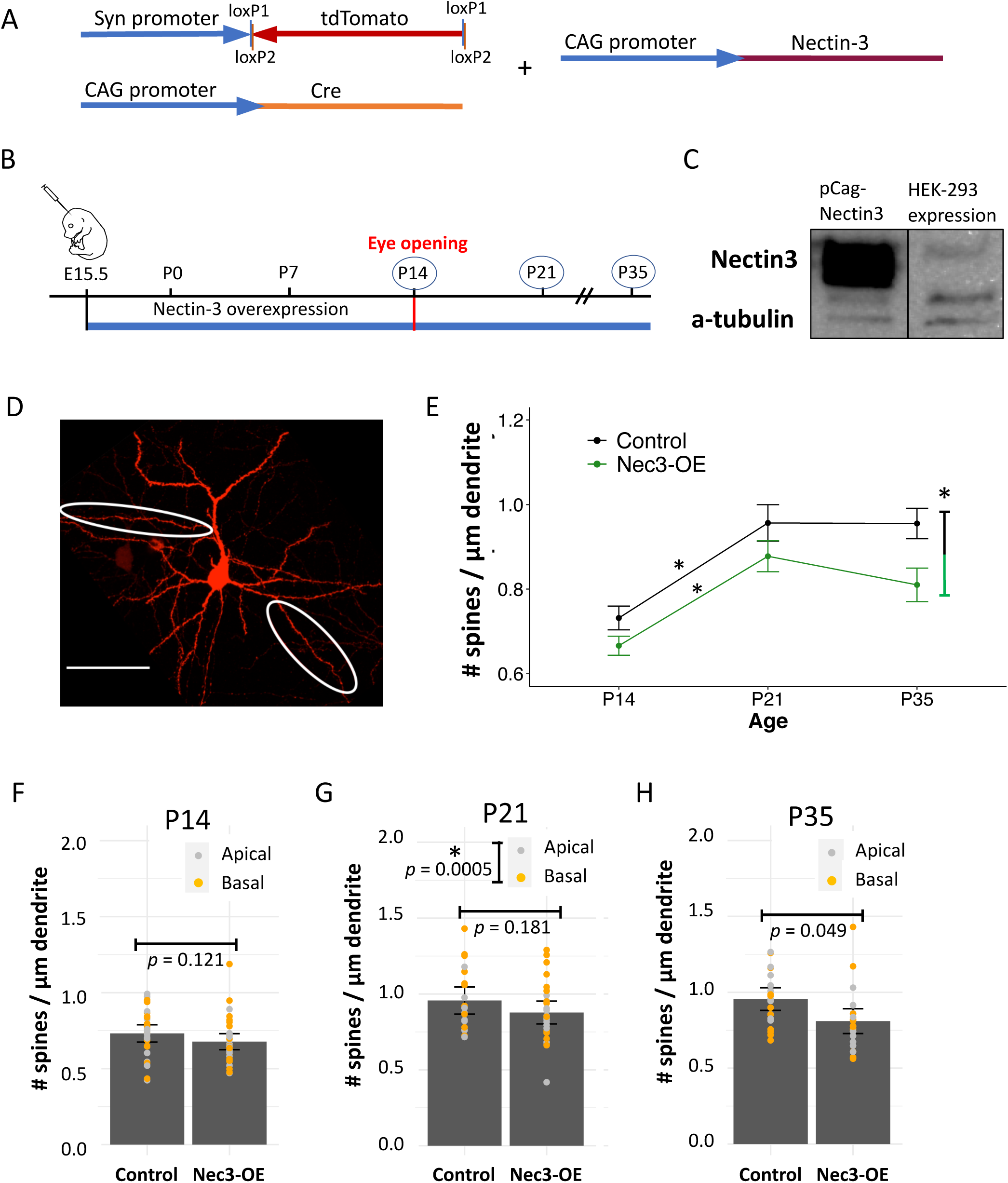
Nectin-3 overexpression beginning at E15.5 decreases dendritic spine density. **(a)** A Nectin-3 expression construct was developed and co-electroporated into in developing L2/3 neurons with a Cre-dependent tdTomato construct and a Cre-expression plasmid. **(b)** Nectin-3 overexpression constructs were introduced to L2/3 cortical neurons by *in utero* electroporation at E15.5. Dendritic spine densities were measured at three time points: P14, P21, and P35. **(c)** Western blot analysis after transfection of HEK-293 cells with a Nectin-3 overexpression vector. Greatly increased Nectin-3 expression is observed relative to endogenous levels in HEK-293 cells. **(d)** Representative image of an electroporated neuron with select apical and basal dendrites circled (scale bar = 50 µm). **(e)** Nectin-3 overexpression produced an overall decrease in dendritic spine densities compared to control (scramble shRNA) neurons (*p* = 0.00437, Bonferroni critical value, *p* < 0.016). As in control neurons, spine densities significantly increased between P14 and P21 (*p* = 0.00017), indicating spinogenesis was intact. Apical and basal dendrites are pooled in this analysis. Significance is denoted by ‘*’, and *p* values are listed in Additional File 1. **(f)** Nec3-OE and control dendritic spine densities at P14. Apical (grey) and basal (orange) dendritic spine densities are plotted along with error bars denoting the standard errors of the mean. **(g)** Nec3-OE and control dendritic spine densities were not different at P21. However, a significant difference in spine densities was observed between apical and basal dendrites at this time point (averaged over condition). **(h)** Dendritic spine densities for Nec3-OE neurons trended lower than control neurons at P35 (*p* = 0.049), but this trend was not significant (Bonferroni critical value, *p* < 0.00192).

**Figure 5:**
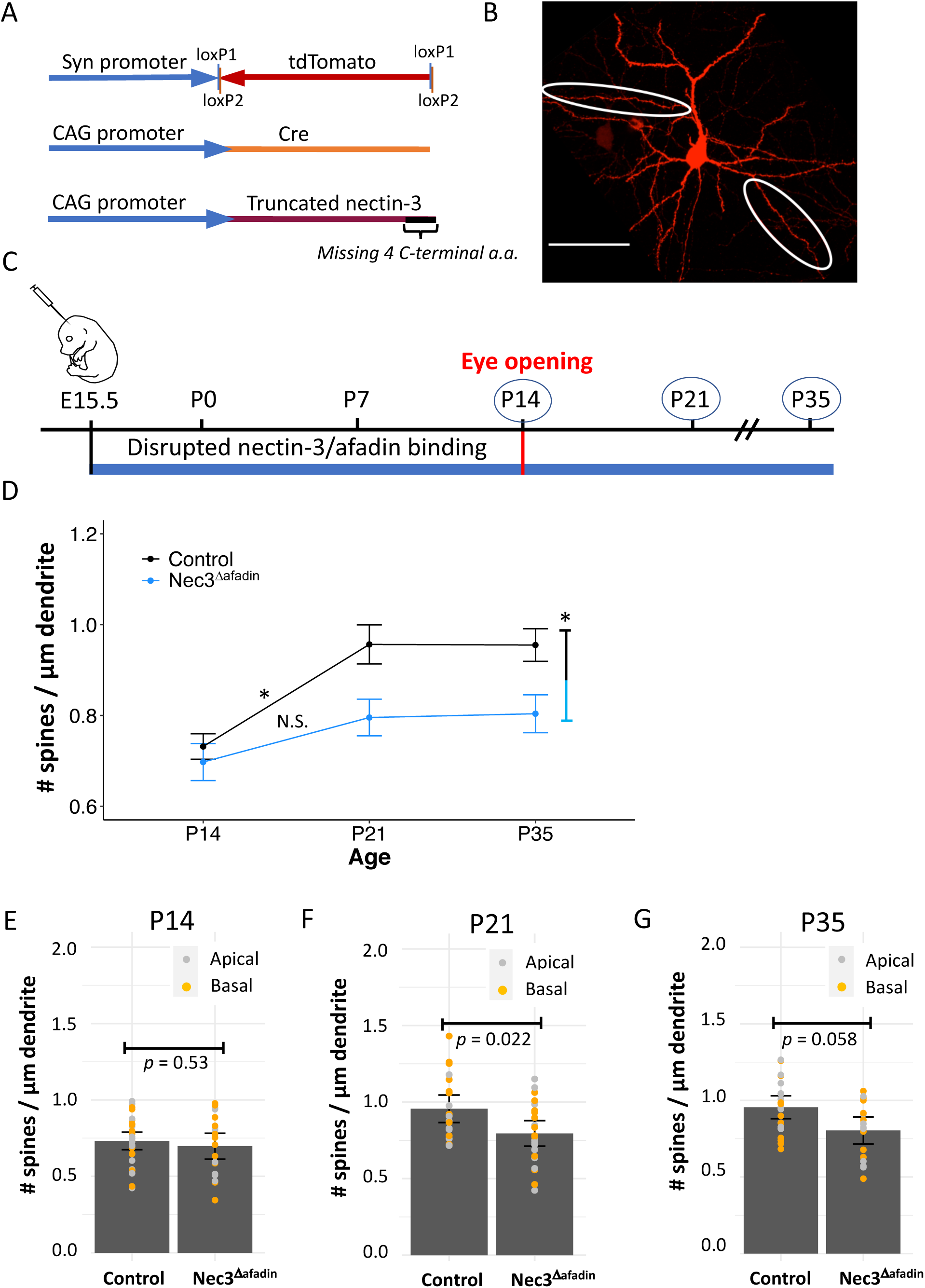
Overexpression of Nec3^Δafadin^ at E15.5 restricts spine formation after eye opening. **(a)** To block the interaction between Nectin-3 and Afadin in developing L2/3 neurons, a construct driving the expression of a truncated Nectin-3 lacking its Afadin binding site was co-electroporated with a Cre-dependent tdTomato construct and a Cre-expression plasmid. **(b)** Representative image of an electroporated neuron with select apical and basal dendrites circled (scale bar = 50 µm). **(c)** A truncated Nectin-3 expression construct, for the dominant negative inhibition of Nectin-3/Afadin binding (Nec3^Δafadin^), was introduced to L2/3 cortical neurons by *in utero* electroporation at E15.5. Dendritic spine densities were measured at P14, P21, or P35. **(d)** Nec3^Δafadin^ produced an overall decrease in dendritic spine densities compared to control (scramble shRNA) neurons when all time points were considered (*p* = 0.0041). In addition, dendritic spine densities did not significantly increase between P14 and P21 (*p* = 0.1765). Apical and basal dendrites are pooled in this analysis. Significance is denoted by ‘*’, and *p* values are listed in Additional File 1. **(e)** Nec3^Δafadin^ and control dendritic spine densities at P14. Apical (grey) and basal (orange) dendritic spine densities are plotted along with error bars denoting the standard errors of the mean. **(f)** Nec3^Δafadin^ and control dendritic spine densities at P21. Dendritic spine densities trended lower at P21 in Nec3^Δafadin^ neurons compared to control (*p* = 0.02215), but this trend was not significant (Bonferroni critical value, *p* < 0.00192). **(g)** Nec3^Δafadin^ and control dendritic spine densities at P35.

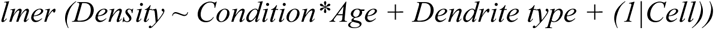

Differences between conditions were tested using an ANOVA, and any significant interactions between ‘condition’ and ‘age’ were noted (Additional File 1). The three comparisons made in this way were considered a family, and a Bonferroni corrected *p* value of 0.016 was used as a cut-off to establish significance (correction for a family of 3 comparisons). *p* values for differences between ages or conditions, as well as between apical and basal dendrite types, are listed for each comparison in Additional File 1.

Finally, we examined apical and basal dendrites independently for developmental changes in dendritic spine densities. For this analysis, we used the ‘car’ package in R to perform an ANOVA where ‘density’ is tested against ‘condition’ and ‘age’ using the following model:

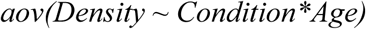

A Tukey’s post-hoc analysis (*TukeyHSD*) was then performed on each model to identify differences between individual experimental groups and select *p* values from this analysis are listed in Additional File 1 (Figure 6). Overall differences in spine densities between ages (all conditions included) and conditions (all ages included) were also examined for apical and basal dendrites separately, and a 95% confidence level was used to establish significance (*p* < 0.05, Additional File 1).

**Figure 6:**
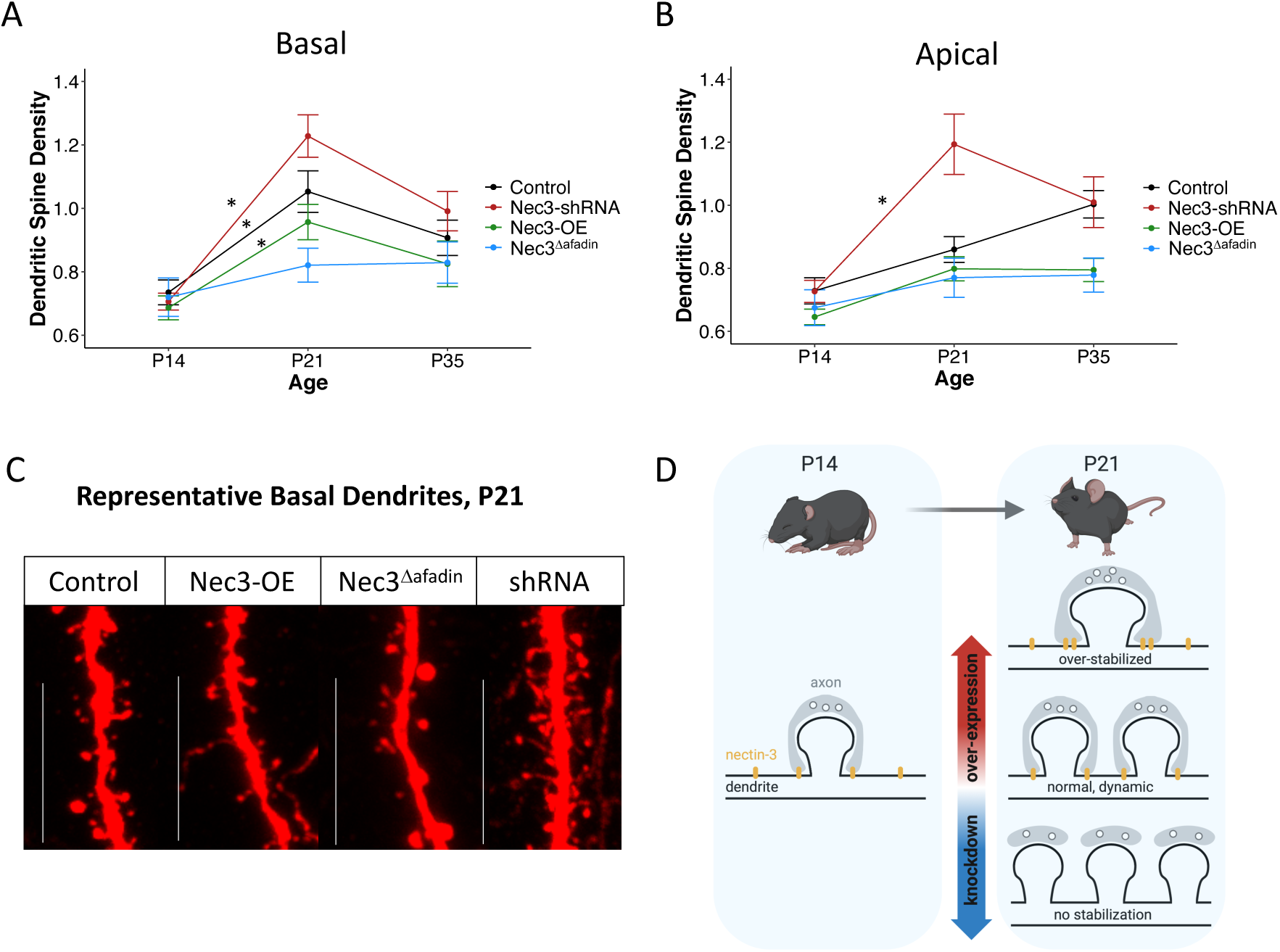
Apical and basal dendritic spine densities respond similarly to Nectin-3 manipulation, **(a)** Developmental trends in dendritic spine densities considering only basal dendrites. All conditions except Nec3^Δafadin^ show significant increases in spine densities between P14 and P21 when considered individually. Significance is denoted by ‘*’, and *p* values are listed in Additional File 1. **(b)** Developmental trends in dendritic spine densities considering only apical dendrites. Only the Nec3-shRNA condition shows a significant increase in spine densities between P14 and P21. Significance is denoted by ‘*’, and *p* values are listed in Additional File 1. **(c)** Images of representative dendrites from neurons electroporated at E15.5 with scramble shRNA (control), full-length Nectin-3 (Nec3), truncated Nectin-3 (Nec3^Δafadin^), or Nectin-3 shRNA (shRNA) expression constructs and assayed at P21 (scale bar = 15 µm). **(d)** Model summarizing results (created with BioRender.com). After eye opening, L2/3 neurons experience an increase in dendritic spine densities. This period of spinogenesis may first require a weakening of synaptic adhesion at existing synapses. In this model, the over-stabilization of existing spines prevents new spine formation, while reduced synaptic adhesion facilitates new spine formation. It is further possible a similar but opposing mechanism facilitates pruning between P21 and P35; the strengthening of select spines by increased cell-adhesion facilitates the removal of neighboring weak spines. As a component of synaptic PAJs, Nectin-3 may facilitate both the accumulation and removal of adhesion molecules at synaptic sites, perhaps through its association with Afadin.

## Results

### Nectin-3 and Nectin-1 are expressed in upper layers of visual cortex

Previous studies have shown a role for Nectin-3 and its binding partner Nectin-1 in the formation and maintenance of hippocampal synapses (Figure 1a) [11,28,31]. To determine the expression pattern of Nectin-3 and Nectin-1 in the developing visual cortex, we used *in situ* hybridization (ISH), and found that both Nectin-3 and Nectin-1 were enriched in layer 2/3 (L2/3) cortical neurons just after eye opening (P16; Figure 1b). This confirmed ISH data found at Allen Brain Atlas, showing enriched L2/3 expression of both nectins beginning in late-embryonic/early-postnatal development and continuing through adulthood [37].

### Nectin-3 knockdown at E15.5 increases spine density at P21

To knockdown Nectin-3 in developing neurons *in vivo*, we first designed new plasmid vectors allowing the Cre-dependent expression of Nectin-3 short-hairpin RNA (shRNA) or scrambled control shRNA (Figure 2a). We used previously published 19 bp siRNA sequences [40] to design our Nectin-3 shRNA oligos and cloned these hairpins into the Cre-dependent pSico vector (Addgene plasmid #11578) [46]. To test for knockdown of Nectin-3, we co-transfected a Nectin-3 expression plasmid with our new Nectin-3 shRNA constructs (with the floxed GFP-Stop sequence removed, see Methods) into HEK-293 cells. Western blot analysis indicated that our newly designed Nectin-3 shRNA constructs effectively reduced Nectin-3 protein expression when compared to cells co-transfected with a scrambled shRNA construct (technical replicates, N = 2, Figure 2b). This is consistent with previously published data showing Nectin-3 was knocked down in cultured cortical neurons using the same siRNA sequences [40].

We used *in utero* electroporation to introduce either or both Nectin-3 shRNA1 and shRNA2 or scrambled shRNA (Figure 2a, b) to developing L2/3 cortical neurons at E15.5 (Figure 2b, c). shRNA constructs were co-electroporated with a Cre-dependent tdTomato construct and a low concentration of a pCag-Cre plasmid (Figure 2a). For this experiment, Cre-negative neurons (expressing the floxed GFP-stop sequence in the pSico construct) fluoresced green, and Cre-positive neurons (excised pSico GFP-stop sequence and recombined FLEX tdTomato construct) fluoresced red (Figure 2d). All cells migrated normally to L2/3 (Figure 2d). We identified tdTomato expressing neurons in L2/3 of primary visual cortex (V1) and assayed both apical and basal dendrites to determine significant changes in dendritic spine densities (Figure 2d, inset). We assayed dendritic spine densities at three developmental time points: eye opening (P14), one week after eye opening (P21), and at the close of the critical period for ocular dominance plasticity (P35; Figure 2c). This allowed us to assess Nectin-3 function during critical periods in the development of visual cortex for synapse formation (P14-P21) [18], as well as synaptic refinement (P21-P35) [1,22,52].

Knocking down Nectin-3 at E15.5 significantly impacted dendritic spine densities during postnatal development. Neurons electroporated with either scrambled shRNA (control) or Nectin-3 shRNA constructs showed an increase in spine densities between P14 and P21, consistent with previous reports of synaptogenesis following eye opening in visual cortex (Figure 2e, significance denoted by ‘*’) [18]. However, Nectin-3 knockdown amplified this change, yielding significantly higher spine densities than control at P21 (Figure 2e, g). By P35, the dendritic spine densities of Nec3-shRNA neurons were no longer significantly different from control neurons. Collectively, these results are consistent with a model where the extended knockdown of Nectin-3 leads to an overproduction of weak spines after eye opening, which are readily pruned over the critical period for ocular dominance plasticity (P21-P35) due to reduced stability.

### Nectin-3 knockdown at ∼P19 increases spine density at P35

Early (E15.5) shRNA knockdown of Nectin-3 yielded an overproduction of dendritic spines after eye opening, significantly increasing dendritic spine densities at P21 relative to control. On the other hand, dendritic spine densities between Nec3-shRNA and control neurons were not different at P35, indicating that the extra spines produced at P21 may have been removed over the critical period for ODP. We next wanted to test whether the initial overproduction of spines at P21 after Nectin-3 knockdown directly led to compensatory spine pruning between P21 and P35. To accomplish this, we selectively knocked down Nectin-3 after the post-eye-opening period of synaptogenesis and assessed dendritic spine densities at P35. For this experiment, we co-electroporated the same Cre-dependent Nec3-shRNA constructs (or control scramble shRNA) and a Cre-dependent tdTomato construct into transgenic CaMKII-Cre mice at E15.5 (Figure 3a, c). In a previous study, CaMKII-Cre driven Cre/loxP recombination was first observed in cortex and hippocampus at P19 and expression was described as robust by P23 [41]. From this, we concluded that the knockdown of Nectin-3 likely occurred in most cells around the start of the critical period for ocular dominance plasticity (P21). Both apical and basal dendritic spine densities were analyzed at P35, to allow time for full shRNA expression and knockdown of Nectin-3 in L2/3 cortical neurons (Figure 3b, c). Unlike early (E15.5) shRNA knockdown of Nectin-3, late Nectin-3 knockdown (∼P19) significantly increased dendritic spine densities at P35 relative to control (Figure 3d). This increase in spine density was not specific to apical or basal dendrites (Figure 3d, Additional File 1). From this we conclude that Nectin-3 may also facilitate pruning over the critical period for ODP.

### Nectin-3 overexpression at E15.5 decreases dendritic spine density

We next examined the effect of overexpressing Nectin-3 in L2/3 cortical neurons over development. We generated a newly designed Nectin-3 expression plasmid, which dramatically increases Nectin-3 expression after transfection into HEK-293 cells (Figure 4a, b). This Nectin-3 overexpression construct was electroporated into developing L2/3 neurons at E15.5, and apical and basal dendritic spine densities were analyzed at P14, P21, and P35 (Figure 4c, d). Dendritic spine densities on both control (scramble shRNA) and Nectin-3 overexpression (Nec3-OE) neurons increased significantly between P14 and P21 (control, P14-P21: *p* = 0.00017; Nec3-OE, P14-P21: *p* = 0.00027, Figure 4e). However, when all time points were considered together, Nec3-OE significantly decreased dendritic spine densities relative to control neurons (*p* = 0.0044) (Figure 4e). The greatest difference between Nec3-OE and control dendritic spine densities was observed at P35. Consistent with our shRNA experiments, these data further indicate that Nectin-3 expression may restrict spinogenesis over development and facilitate pruning over the critical period for ODP (P21 – P35).

Our examination of combined Nec3-OE and control datasets also revealed a potential difference in the degree of spinogenesis experienced by apical and basal dendrites after eye opening. For every comparison between experimental groups, we used a linear mixed model which included either ‘condition’ or ‘age’ as well as ‘dendrite type’ (apical vs. basal) as fixed variables (see methods for a detailed description of statistical approach). As such, every comparison was tested by ANOVA for differences in dendritic spine densities both between experimental groups as well as between apical and basal dendrites (averaged over experimental groups). Apical and basal dendritic spine densities trended different at P21 for both Nec3-OE and control conditions when tested independently, though our study failed to demonstrate statistical significance using a Bonferroni critical value of 0.0036 (t-test, Additional File 1, Metadata). When these datasets are combined, however, significant differences between Additional File 1, Figure 4g). It is important to note that no difference between Nec3-OE and control conditions was observed at P21 (Figure 4g), indicating that the overexpression of Nectin-3 does not significantly impact dendritic spine densities at this time point. This result does suggest, however, that the apical dendrites sampled in this study (secondary proximal dendrites) may not increase their spine densities after eye opening to the same degree as basal dendrites under the conditions of normal or above normal Nectin-3 expression (Figure 4g, Figure 6). From this we conclude that dendrite type may also be an important consideration when evaluating developmental changes in dendritic spine densities following eye opening, as discussed below.

### Nectin-3 binding to Afadin may limit Nectin-induced reduction in spine density

Nectin proteins consist of a three extracellular Ig domains, a transmembrane fragment, and multiple intracellular C-terminal domains with distinct and independent functions [11,39,53–55]. Most nectins, including Nectin-3, have a conserved motif of four amino acids at their cytoplasmic tail that binds the PDZ domain of Afadin [11,28,38,39]. Nectin-3/Afadin binding is required for the interaction of Nectin-3 with the actin cytoskeleton and the organization of PAJs in cooperation with N-cadherin (Figure 1a) [11,26–30,56]. Nectins can also recruit and activate c-Src leading to the downstream activation of Cdc42 and Rac, which facilitate the formation of filopodia and lamellipodia, respectively [54,57–59]. The Nectin-1 C-terminus, but not the Afadin binding site, was necessary for the activation of Cdc42 and Rac [57]. The extracellular domain of Nectin-3 has also been shown to act independently of its C-terminus to bind Nectin-1, leading to Nectin-1 intracellular signaling and cadherin recruitment [54,57,60]. It is unclear which of Nectin-3’s multiple functions may regulate dendritic spine formation during critical periods of postnatal development.

We next tested whether the observed Nectin-3-induced reduction in spine density required an interaction between Nectin-3 and Afadin. To accomplish this, we used *in utero* electroporation to overexpress a Nectin-3 protein lacking the four C-terminal amino acids required to bind Afadin (Nec3^Δafadin^) [11,28,38,39]. Developing L2/3 neurons were electroporated with Nec3^Δafadin^ at E15.5, and dendritic spine densities were assayed at P14, P21, and P35 (Figure 5b, c). It should be noted that the Nec3^Δafadin^ protein can still bind Nectin-1 and activate Cdc42 and Rac signaling [54,57–59]. Similar to the Nec3-OE condition, Nec3^Δafadin^ produced an overall decrease in dendritic spine densities relative to control neurons (scramble shRNA) (Figure 5d). From this we conclude that Nectin-3 utilizes an Afadin-independent pathway to restrict spine formation during this developmental period. Unlike Nec3-OE, the greatest difference between Nec3^Δafadin^ and control neurons was observed one week following eye opening, at P21. Furthermore, we observed a significant developmental increase in dendritic spine densities between P14 and P21 in control and Nec3-OE conditions, but not Nec3^Δafadin^ (Figure 5d, 4e). From this we conclude that, in the condition of high Nectin-3 levels, the interaction between Nectin-3 and Afadin may facilitate normal spine formation following eye opening. Collectively, these results support a model whereby spine formation after eye opening is restricted by the extracellular adhesive properties of Nectin-3 but facilitated by a Nectin-3-Afadin interaction, which could function to redistribute Nectin-3 away from synaptic sites at this time.

### Apical and basal dendritic spine densities respond similarly to Nectin-3 manipulation, despite demonstrating different patterns of developmental change

We next examined apical and basal dendrite types separately to identify potential differences in the way dendritic spine densities change over development. There are previous reports of neurons experiencing a period of intense synaptogenesis following eye opening (mouse P14–P21), followed by the critical period for optical dominance plasticity (mouse P21–P35), which is associated with synaptic refinement and pruning [18–20,22,61,62]. When pooling apical and basal dendrites (Figures 2, 4, 5), or considering basal dendrites alone (Figure 6a), all conditions except Nec3^Δafadin^ demonstrated a significant increase in dendritic spine densities from P14 to P21. Furthermore, the largest differences in dendritic spine densities between conditions was observed at P21 (Figure 6a, c, Additional File 2: Figure S1). This is consistent with previous studies examining synapse formation at this time, and indicates that Nectin-3 may have a role in regulating synaptogenesis following eye opening [18]. In addition, when basal dendrites are considered alone, dendritic spine densities trend downwards between P21 and P35 for all conditions except Nec3^Δafadin^. While none of these trends are statistically significant when considered individually, a significant decrease in spine densities is observed when all conditions are considered together (Tukey’s post-hoc test, P21–P35, *p* = 0.022, Additional File 1). This result indicates synapse pruning may be occurring on basal dendrites between P21 and P35, though we acknowledge that the manipulation of Nectin-3 expression may have affected this result (Figure 6a). We conclude that our observed developmental changes in basal dendritic spine densities are consistent with previous studies reporting increases in dendritic spine and synapse number following eye opening and synapse pruning over the critical period for ODP [2,4,18–22].

The apical dendrites sampled in this study may form dendritic spines at a different rate than basal dendrites after eye opening. Unlike basal dendrites, when apical dendrites are considered alone (Figure 6b), control and Nec3-OE dendritic spine densities do not significantly change between P14 and P21 (Figure 6b). Furthermore, dendritic spine densities continue to increase between P21 and P35 in the control condition alone, demonstrating a significant difference in spine density at P35 relative to P14 (*p* = 0.0058, Additional File 1). This indicates that, unlike other dendrite types [22,63], the apical dendrites we sampled here (proximal secondary apical dendrites, distinct from apical tufts) may be slower to increase dendritic spine densities after eye opening and may continue to increase spine densities over the critical period for ODP. Interestingly, overexpressing Nectin-3 both with and without its Afadin binding site (Nec3-OE and Nec3^Δafadin^) may prevent this further increase in apical spine densities between P21 and P35 (no significant differences between time points observed, Additional File 1). This further supports a model whereby increased adhesion caused by high expression of Nectin-3 is detrimental to normal developmental increases in dendritic spine densities (Figure 6d). We conclude that the observed differences between apical and basal dendrites do not fundamentally change the outcome of these Nectin-3 manipulations: decreasing Nectin-3 caused increased spine densities, and overexpressing Nectin-3, regardless of whether the protein had an intact Afadin binding site, reduced spine densities (Figure 6d).

## Discussion

During the first three weeks following eye opening, the connectivity and function of murine visual cortex changes rapidly to adapt to the onset of visually evoked neuronal activity [2,18,62,64]. This reorganization of synaptic connections is thought to occur over two distinct critical periods, P14-P21 (eye opening), and P21-P35 (the critical period for ODP) [2,4,18–20,22]. It is largely unknown how cell adhesion molecules, including the L2/3 enriched binding partners Nectin-3 and Nectin-1, may affect synapse formation or pruning during these critical periods of visual cortex development. Previous studies examining hippocampal neurons have identified different roles for Nectin-3 in regulating dendritic spine densities depending on age and system (*in vivo* vs. *in vitro*) [11,34,36]. Two studies found dendritic spine densities decreased with Nectin-3 knockdown in adult mouse hippocampus *in vivo* [34,36]. On the other hand, blockade of Nectin-1 to Nectin-3 binding in developing hippocampal neurons *in vitro* increased dendritic spine densities and decreased spine size [11]. Consistent with the latter, here we identify a potential function for Nectin-3 in limiting dendritic spine densities following eye opening in L2/3 visual cortex. These observations collectively indicate that Nectin-3 may have a role in synapse formation, pruning, or maintenance that changes throughout development and aging.

To identify potential roles for Nectin-3 in postnatal cortical development, we used the powerful technique, *in utero* electroporation, to knockdown (Nectin-3 shRNA) or overexpress (Nec3-OE) Nectin-3 in developing L2/3 visual cortical neurons *in vivo*. We additionally overexpressed a Nectin-3 lacking its Afadin binding site (Nec3^Δafadin^), which should have acted as a dominant negative for Nectin-3/Afadin binding, overwhelming endogenous Nectin-3 function [40,65]. We then assayed dendritic spine densities on neurons with modified Nectin-3 expression at critical ages in the development of visual cortex (P14, P21 and P35). Consistent with previous literature [18,19,21], we observed a significant increase in dendritic spine densities between P14 and P21 for all conditions except Nec3^Δafadin^. In control neurons, this increase in spine densities occurred rapidly on basal dendrites between P14 and P21, but was more prolonged for apical dendrites, showing a significant increase between P14 and P35, but not between P14 and P21 (Figure 6, Additional File 1). Nec3-shRNA increased dendritic spine densities at P21, indicating that this treatment may have ‘lifted the breaks’ on spine formation at this time. On the other hand, Nec3-OE and Nec3^Δafadin^ both yielded significant overall decreases in dendritic spine densities, which were most pronounced following periods of spine formation for each dendrite type (P21 for basal dendrites and P35 for apical dendrites, Figure 6). Nec3^Δafadin^ was the only condition to prevent a significant increase in basal (or overall combined) dendritic spine densities between P14 and P21 (Figure 4, 6a). Taken together, these results indicate that Nectin-3 may function to restrict dendritic spine formation after eye opening, and that Nectin-3/Afadin binding may act to counterbalance this function.

One possibility is that Nectin-3 may act to restrict the homeostatic scaling of synaptic strength in L2/3 neurons following eye opening in V1. It was previously shown that the frequency of spontaneous miniature excitatory synaptic currents (mEPSCs, an indicator of synapse number) increases in upper layer cortical neurons after eye opening [18]. This occurs simultaneously with a decrease in mEPSC amplitude, an indicator of synaptic strength [18]. Since synapse number and visual input both increase after eye opening, a decrease in synapse strength may be necessary to prevent neuronal overexcitability (homeostatic scaling) [18,66]. If, by extension, high synaptic strength is prohibitive to increases in synapse number, Nectin-3 knockdown may facilitate increases in synapse number by decreasing synaptic strength. It has previously been suggested that decreased synaptic adhesion may increase synapse number either to compensate for reduced synaptic strength [11], or by allowing the redistribution of membrane components to sites where new synapses form [67]. Another possibility is that Nectin-3 knockdown facilitates new spine formation by freeing up synaptic molecules which would otherwise be restricted by PAJs (Figure 1). In this way, Nectin-3 and other synaptic adhesion molecules may help neurons balance synapse strength and number following eye opening.

Our study indicates that the association of Nectin-3 with Afadin, and by extension the actin cytoskeleton, may facilitate spine formation after eye opening. Cytoskeletal components have previously been shown to both influence and be influenced by the cell-adhesion molecules they associate with [68], and nectin proteins appear to be no exception [30]. In immature (7 days in vitro (DIV)) developing cultured hippocampal neurons, actin depolymerization (latrunculin A treatment) prevented the localization of Nectin-1 to synaptic sites and decreased the size of synaptophysin clusters (an indicator of reduced synaptic strength) [30]. In older neurons (21 DIV), actin depolymerization increased the density of synaptophysin clusters (an indicator of increased synapse number) and decreased the size of Nectin-1 puncta at synaptic sites [30]. Accordingly, it is possible that cytoskeletal changes following eye opening [20] may also influence the position and function of Nectin-3, perhaps facilitating its removal from synaptic sites. Further studies are necessary to determine whether the redistribution of Nectin-3 away from synaptic sites may have a role in homeostatic synaptic scaling and synapse formation after eye opening [18].

While eye opening is associated with an increase in synapse number and a decrease in synapse strength, the opposite is true during critical periods for experience-dependent synaptic plasticity (critical period for ODP in V1, P21–P35) [61,69]. At these times, active synapses require a mechanism for selective strengthening while inactive synapses are weakened and pruned [61,69]. This may be facilitated by the selective accumulation of synaptic adhesion molecules at active synapses and competitive removal from inactive synapses [7,9,17]. For example, it was previously demonstrated that competition between dendritic spines for N-cadherin, which is colocalized with nectins at synapses, may drive experience-dependent pruning in mouse somatosensory cortex [9,11]. Here we show that when Nectin-3 is knocked down acutely (∼P19–P35) during the critical period for ODP, using CaMKII-Cre transgenic mice [41], spine densities are higher at P35 when compared to control neurons (Figure 3). This indicates that Nectin-3 may facilitate synapse pruning during the critical period for ODP, though it is possible that synapse formation was also affected by this manipulation. One hypothesis is that competition between spines for Nectin-3 and its associated molecules may lead to the selective strengthening of active spines and weakening of inactive spines [9]. Several molecules associated with Nectin-3 at PAJs, including N-cadherin, catenins, and Afadin, have been shown to localize to synapses in response to neuronal activity (Figure 1a) [7,8,70–75]. Interestingly, Nectin-3 has been shown to recruit N-cadherin (C-terminal domain) to adherens junctions in an Afadin-dependent manner [53]. Further experiments are necessary to determine whether Nectin-3 may also experience activity-dependent synaptic localization or may influence the localization of associated PAJ molecules. In this way, Nectin-3 could influence the activity-dependent refinement of synaptic connections between L2/3 cortical neurons during the critical period for ocular dominance plasticity in V1.

There are a number of challenges associated with using *in utero* electroporation to manipulate protein expression in developing neurons. Plasmid concentration in a given electroporated neuron can vary greatly, potentially leading to increased phenotypic variability from neuron to neuron within a single condition. In addition, there is high endogenous variability in spine densities between dendrites and cells in L2/3 of visual cortex. We focused on proximal apical and basal dendrites beginning one branch away from the soma (secondary dendrites) in order to limit the effect of this variability, but it was still difficult to obtain robust estimates of population differences. Previous studies of developing mouse or rat visual cortex have varied in their descriptions of dendritic spine densities over the critical period for ODP [19,20,22,76], and different dendrite types on L3 pyramidal neurons have been shown to experience different degrees of pruning [19]. Our results indicate that apical and basal (secondary proximal) dendritic spine densities may experience different developmental trajectories after eye opening. However, most overall effects, including increased spine densities with Nectin-3 knockdown and decreased spine densities with Nectin-3 overexpression, were consistent between apical and basal dendrites. For this reason, we decided to combine apical and basal dendrites for most of our analyses, and we consider dendrite type (apical vs basal) as a fixed effect in our statistical models (see methods). We further believe combining both dendrite types provided a more accurate approximation of the overall cellular effect of Nectin-3 manipulation.

In this study, we identify Nectin-3 dependent changes to dendritic spine densities on L2/3 visual cortical neurons that become evident after eye opening and are dependent on time of knockdown and association with Afadin. We hypothesize that Nectin-3 may have roles in the formation and strengthening specific synapses; preventing runaway spine formation at P21 and facilitating the selective removal of inactive spines between P21 and P35. Further experiments are necessary to validate these hypothesized roles for Nectin-3 in the development of L2/3 visual cortical neurons. Similar to the experiment performed by Bian et al. (2015), it would be informative to examine whether Nectin-1 to Nectin-3 binding at specific synapses in cultured neurons facilitates the competitive accumulation of cadherin/catenin complexes to those synapses and reduction at other synapses [9]. Electrophysiological recordings or calcium indicator imaging of neuronal activity could help determine whether Nectin-3 expression facilitates the refinement of visual response properties (such as orientation selectivity, receptive field structure, or contextual modulation) in L2/3 cortical neurons, which would be expected if Nectin-3 enables selective synaptic strengthening and pruning. Furthermore, examining Nectin-3-dependent effects on spine density and synapse function after dark rearing could help address whether Nectin-3 is functionally regulated by neuronal activity. In conclusion, our demonstration of a novel mechanism for regulating spine formation after eye opening should provide a foundation for future studies of the role of nectins in the assembly of cortical circuits.

## Conclusions

We conclude that the expression level of Nectin-3 in layer 2/3 visual cortical neurons influences dendritic spine densities after eye opening. We suggest that tightly regulating Nectin-3 at synapses, perhaps through an interaction with Afadin, may provide a mechanism for controlling the strength and number of dendritic spines on specific neurons during critical periods of cortical development.

## Additional Files

### Additional File 1

Summary of all statistical analyses and *p* values for comparisons between condition, ages, and dendrite type. The type of analysis as well as the Bonferroni critical value used to establish significance, are indicated. Whether the data was first transformed to correct for non-normality or heteroskedasticity is also indicated, and final *p* values for Shapiro and Levine tests (after any necessary data transformations) are shown. An additional sheet labeled ‘Metadata’ documents the number of animals and dendrites sampled for each condition. This sheet also includes *p* values describing individual t-test comparisons of apical and basal dendrites for each condition. (.xlsx file, 20 KB)

### Additional File 2: Figure S1

Nectin-3 overexpression and loss of function most significantly affects dendritic spine densities at P21. (**a**) No significant differences in dendritic spine densities are observed at P14 between neurons electroporated at E15.5 with control, Nec3-shRNA, Nec3-OE, or Nec3^Δafadin^ constructs. Individual counts for apical and basal dendrites are shown in grey and black, respectively. (**b**) Significant in dendritic spine densities emerge at P21between neurons electroporated at E15.5 with control, Nec3-shRNA, Nec3-OE, or Nec3^Δafadin^ constructs. (**c**) No significant differences in dendritic spine densities are observed at P35 between neurons electroporated at E15.5 with control, Nec3-shRNA, Nec3-OE, or Nec3^Δafadin^ constructs. (.pdf file, 134 KB)

## List of abbreviations

PAJ: Puncta Adherentia Junction
Nec3-shRNA: Nectin-3 short hairpin RNA
Nec3-OE: Nectin-3 overexpression
Nec3^Δafadin^: Nectin-3 missing the Afadin binding site
siRNA: small interfering RNA
V1: Primary visual cortex
L2/3: Layer 2/3
P14: Postnatal day 14
ODP: Ocular dominance plasticity
IHC: immunohistochemistry.

## Declarations

## Acknowledgements

The authors are grateful to Mandi Severson for her extensive work imaging dendrites and processing tissue for histological analysis. We would also like to acknowledge Emily Hill, Brynna Paros, Claire (Hoa) Bui, and Nathan Nguyen for their hard work processing tissue and counting dendritic spines. We are also very grateful to Judit Pungor for helping design the schematic for Figure 1a; Nick Sattler and Rachel Lukowicz for their contributions to the project during their rotations in the Niell lab; and Bree Mohr for performing western blot analysis to determine Nectin-3 knockdown.

## Authors’ contributions

Electroporation experiments in Figures 2-6 were performed by JT and PRLP. Experiments in Figure 1, molecular cloning and construct design, and all data and statistical analyses were performed by JT. The manuscript was written by JT and edited by PRLP, CQD, and CMN. All authors read and approved of the final manuscript.

## References

1. Katz LC, Shatz CJ. Synaptic activity and the construction of cortical circuits. Science (80-.). 1996. p. 1133–8.

2. Chen C-C, Lu J, Zuo Y. Spatiotemporal dynamics of dendritic spines in the living brain. Front Neuroanat. Frontiers; 2014;8:28.

3. Hackett TA, Guo Y, Clause A, Hackett NJ, Garbett K, Zhang P, et al. Transcriptional maturation of the mouse auditory forebrain. BMC Genomics. BioMed Central; 2015;16:606.

4. Benoit J, Ayoub AE, Rakic P. Transcriptomics of critical period of visual cortical plasticity in mice. Proc Natl Acad Sci U S A. National Academy of Sciences; 2015;112:8094–9.

5. Tomorsky J, DeBlander L, Kentros CG, Doe CQ, Niell CM. TU-Tagging: A Method for Identifying Layer-Enriched Neuronal Genes in Developing Mouse Visual Cortex. eNeuro. Society for Neuroscience; 2017;4.

6. Dalva MB, McClelland AC, Kayser MS. Cell adhesion molecules: signalling functions at the synapse. Nat Rev Neurosci. Nature Publishing Group; 2007;8:206–20.

7. Xie Z, Photowala H, Cahill ME, Srivastava DP, Woolfrey KM, Shum CY, et al. Coordination of synaptic adhesion with dendritic spine remodeling by aF-6 and kalirin-7. J Neurosci. Society for Neuroscience; 2008;28:6079–91.

8. Bozdagi O, Shan W, Tanaka H, Benson DL, Huntley GW. Increasing numbers of synaptic puncta during late-phase LTP: N-cadherin is synthesized, recruited to synaptic sites, and required for potentiation. Neuron. Cell Press; 2000;28:245–59.

9. Bian WJ, Miao WY, He SJ, Qiu Z, Yu X. Coordinated Spine Pruning and Maturation Mediated by Inter-Spine Competition for Cadherin/Catenin Complexes. Cell. Elsevier; 2015;162:808–22.

10. Takai Y, Shimizu K, Ohtsuka T. The roles of cadherins and nectins in interneuronal synapse formation. Curr. Opin. Neurobiol. Elsevier Ltd; 2003. p. 520–6.

11. Mizoguchi A, Nakanishi H, Kimura K, Matsubara K, Ozaki-Kuroda K, Katata T, et al. Nectin: an adhesion molecule involved in formation of synapses. J Cell Biol. Rockefeller University Press; 2002;156:555–65.

12. Yuste R, Bonhoeffer T. Morphological Changes in Dendritic Spines Associated with Long-Term Synaptic Plasticity. Annu Rev Neurosci. Annual Reviews; 2001;24:1071–89.

13. Trachtenberg JT, Chen BE, Knott GW, Feng G, Sanes JR, Welker E, et al. Long-term in vivo imaging of experience-dependent synaptic plasticity in adult cortex. Nature. 2002;420:788–94.

14. Lin YC, Koleske AJ. Mechanisms of synapse and dendrite maintenance and their disruption in psychiatric and neurodegenerative disorders. Annu Rev Neurosci. NIH Public Access; 2010;33:349–78.

15. Harris KM. Structure, development, and plasticity of dendritic spines. Curr Opin Neurobiol. Elsevier Current Trends; 1999;9:343–8.

16. Arellano JI. Ultrastructure of dendritic spines: correlation between synaptic and spine morphologies. Front Neurosci. Frontiers Media SA; 2007;1:131–43.

17. El-Boustani S, Ip JPK, Breton-Provencher V, Knott GW, Okuno H, Bito H, et al. Locally coordinated synaptic plasticity of visual cortex neurons in vivo. Science. American Association for the Advancement of Science; 2018;360:1349–54.

18. Desai NS, Cudmore RH, Nelson SB, Turrigiano GG. Critical periods for experience-dependent synaptic scaling in visual cortex. Nat Neurosci. 2002;5:783–9.

19. Juraska JM. The development of pyramidal neurons after eye opening in the visual cortex of hooded rats: A quantitative study. J Comp Neurol. 1982;212:208–13.

20. Chen MQ, Bi AL, Zhang YY, Yan Q, Sun YY, Zhang XY, et al. Different patterns of changes between actin dynamics and synaptic density in the rat’s primary visual cortex during a special period of visual development. Brain Res Bull. Elsevier; 2017;132:199–203.

21. Gandhi SP, Cang J, Stryker MP. An eye-opening experience. Nat. Neurosci. Nature Publishing Group; 2005. p. 9–10.

22. Vidal GS, Djurisic M, Brown K, Sapp RW, Shatz CJ. Cell-Autonomous Regulation of Dendritic Spine Density by PirB. eNeuro. Society for Neuroscience; 2016;3.

23. Lendvai B, Stern EA, Chen B, Svoboda K. Experience-dependent plasticity of dendritic spines in the developing rat barrel cortex in vivo. Nature. 2000;404:876–81.

24. Ma L, Qiao Q, Tsai JW, Yang G, Li W, Gan WB. Experience-dependent plasticity of dendritic spines of layer 2/3 pyramidal neurons in the mouse cortex. Dev Neurobiol. NIH Public Access; 2016;76:277–86.

25. Hoy JL, Niell CM. Layer-specific refinement of visual cortex function after eye opening in the awake mouse. J Neurosci. 2015;35:3370–83.

26. Honda T, Sakisaka T, Yamada T, Kumazawa N, Hoshino T, Kajita M, et al. Involvement of nectins in the formation of puncta adherentia junctions and the mossy fiber trajectory in the mouse hippocampus. Mol Cell Neurosci. Academic Press; 2006;31:315–25.

27. Rikitake Y, Mandai K, Takai Y. The role of nectins in different types of cell-cell adhesion. J Cell Sci. 2012;125:3713–22.

28. Takai Y, Nakanishi H. Nectin and afadin: novel organizers of intercellular junctions. J Cell Sci. The Company of Biologists Ltd; 2003;116:17–27.

29. Tachibana K, Nakanishi H, Mandai K, Ozaki K, Ikeda W, Yamamoto Y, et al. Two cell adhesion molecules, nectin and cadherin, interact through their cytoplasmic domain-associated proteins. J Cell Biol. Rockefeller University Press; 2000;150:1161–76.

30. Lim ST, Lim KC, Giuliano RE, Federoff HJ. Temporal and spatial localization of nectin-1 and l-afadin during synaptogenesis in hippocampal neurons. J Comp Neurol. Wiley-Blackwell; 2008;507:1228–44.

31. Honda T, Sakisaka T, Yamada T, Kumazawa N, Hoshino T, Kajita M, et al. Involvement of nectins in the formation of puncta adherentia junctions and the mossy fiber trajectory in the mouse hippocampus. Mol Cell Neurosci. Academic Press; 2006;31:315–25.

32. Maurin H, Seymour CM, Lechat B, Borghgraef P, Devijver H, Jaworski T, et al. Tauopathy Differentially Affects Cell Adhesion Molecules in Mouse Brain: Early Down-Regulation of Nectin-3 in Stratum Lacunosum Moleculare. Vitorica J, editor. PLoS One. 2013;8:e63589.

33. van der Kooij MA, Fantin M, Rejmak E, Grosse J, Zanoletti O, Fournier C, et al. Role for MMP-9 in stress-induced downregulation of nectin-3 in hippocampal CA1 and associated behavioural alterations. Nat Commun. Nature Publishing Group; 2014;5:4995.

34. Wang XX, Li JT, Xie XM, Gu Y, Si TM, Schmidt MV, et al. Nectin-3 modulates the structural plasticity of dentate granule cells and long-term memory. Transl Psychiatry. Nature Publishing Group; 2017;7:e1228.

35. Gong Q, Su YA, Wu C, Si TM, Deussing JM, Schmidt MV., et al. Chronic Stress Reduces Nectin-1 mRNA Levels and Disrupts Dendritic Spine Plasticity in the Adult Mouse Perirhinal Cortex. Front Cell Neurosci. 2018;12:67.

36. Wang XD, Su YA, Wagner KV, Avrabos C, Scharf SH, Hartmann J, et al. Nectin-3 links CRHR1 signaling to stress-induced memory deficits and spine loss. Nat Neurosci. Nature Publishing Group; 2013;16:706–13.

37. © 2008 Allen Institute for Brain Science. Allen Developing Mouse Brain Atlas. Available from: developingmouse.brain-map.org

38. Fujiwara Y, Goda N, Tamashiro T, Narita H, Satomura K, Tenno T, et al. Crystal structure of afadin PDZ domain-nectin-3 complex shows the structural plasticity of the ligand-binding site. Protein Sci. Blackwell Publishing Ltd; 2015;24:376–85.

39. Satoh-Horikawa K, Nakanishi H, Takahashi K, Miyahara M, Nishimura M, Tachibana K, et al. Nectin-3, a new member of immunoglobulin-like cell adhesion molecules that shows homophilic and heterophilic cell-cell adhesion activities. J Biol Chem. American Society for Biochemistry and Molecular Biology; 2000;275:10291–9.

40. Gil-Sanz C, Franco SJ, Martinez-Garay I, Espinosa A, Harkins-Perry S, Müller U. Cajal-Retzius Cells Instruct Neuronal Migration by Coincidence Signaling between Secreted and Contact-Dependent Guidance Cues. Neuron. 2013;79:461–77.

41. Tsien JZ, Chen DF, Gerber D, Tom C, Mercer EH, Anderson DJ, et al. Subregion- and cell type-restricted gene knockout in mouse brain. Cell. Cell Press; 1996;87:1317–26.

42. Oh SW, Harris JA, Ng L, Winslow B, Cain N, Mihalas S, et al. A mesoscale connectome of the mouse brain. Nature. 2014;508:207–14.

43. Wehr M, Hostick U, Kyweriga M, Tan A, Weible AP, Wu H, et al. Transgenic silencing of neurons in the mammalian brain by expression of the allatostatin receptor (AlstR). J Neurophysiol. 2009;102:2554–62.

44. Lein ES, Hawrylycz MJ, Ao N, Ayres M, Bensinger A, Bernard A, et al. Genome-wide atlas of gene expression in the adult mouse brain. Nature. 2007;445:168–76.

45. © 2004 Allen Institute for Brain Science. Allen Mouse Brain Atlas. Available from: mouse.brain-map.org

46. Ventura A, Meissner A, Dillon CP, McManus M, Sharp PA, Van Parijs L, et al. Cre-lox-regulated conditional RNA interference from transgenes. Proc Natl Acad Sci U S A. National Academy of Sciences; 2004;101:10380–5.

47. Mahmood T, Yang PC. Western blot: Technique, theory, and trouble shooting. N Am J Med Sci. 2012;4:429–34.

48. Harwell CC, Parker PRL, Gee SM, Okada A, McConnell SK, Kreitzer AC, et al. Sonic Hedgehog Expression in Corticofugal Projection Neurons Directs Cortical Microcircuit Formation. Neuron. Cell Press; 2012;73:1116–26.

49. Walantus W, Castaneda D, Elias L, Kriegstein A. In utero intraventricular injection and electroporation of E15 mouse embryos. J Vis Exp. 2007.

50. Piscopo DM, Weible AP, Rothbart MK, Posner MI, Niell CM. Changes in white matter in mice resulting from low-frequency brain stimulation. Proc Natl Acad Sci U S A. National Academy of Sciences; 2018;115:E6339–46.

51. Paxinos G, Franklin K. Paxinos and Franklin’s the Mouse Brain in Stereotaxic Coordinates, Fourth Edition. Academic Press; 2013.

52. Espinosa JS, Stryker MP. Development and plasticity of the primary visual cortex. Neuron. 2012;75:230–49.

53. Tanaka Y, Nakanishi H, Kakunaga S, Okabe N, Kawakatsu T, Shimizu K, et al. Role of nectin in formation of E-cadherin-based adherens junctions in keratinocytes: Analysis with the N-cadherin dominant negative mutant. Mol Biol Cell. 2003;14:1597–609.

54. Ogita H, Takai Y. Activation of Rap1, Cdc42, and Rac by Nectin Adhesion System. Methods Enzymol. 2006. p. 415–24.

55. Sakamoto Y, Ogita H, Hirota T, Kawakatsu T, Fukuyama T, Yasumi M, et al. Interaction of integrin alpha(v)beta3 with nectin. Implication in cross-talk between cell-matrix and cell-cell junctions. J Biol Chem. American Society for Biochemistry and Molecular Biology; 2006;281:19631–44.

56. Togashi H, Miyoshi J, Honda T, Sakisaka T, Takai Y, Takeichi M. Interneurite affinity is regulated by heterophilic nectin interactions in concert with the cadherin machinery. J Cell Biol. Rockefeller University Press; 2006;174:141–51.

57. Kawakatsu T, Shimizu K, Honda T, Fukuhara T, Hoshino T, Takai Y. trans-interactions of nectins induce formation of filopodia and lamellipodia through the respective activation of Cdc42 and Rac small G proteins. J Biol Chem. American Society for Biochemistry and Molecular Biology; 2002;277:50749–55.

58. Kawakatsu T, Ogita H, Fukuhara T, Fukuyama T, Minami Y, Shimizu K, et al. Vav2 as a Rac-GDP/GTP exchange factor responsible for the nectin-induced, c-Src- and Cdc42-mediated activation of Rac. J Biol Chem. 2005;280:4940–7.

59. Fukuhara T, Shimizu K, Kawakatsu T, Fukuyama T, Minami Y, Honda T, et al. Activation of Cdc42 by trans interactions of the cell adhesion molecules nectins through c-Src and Cdc42-GEF FRG. J Cell Biol. The Rockefeller University Press; 2004;166:393–405.

60. Honda T, Shimizu K, Kawakatsu T, Yasumi M, Shingai T, Fukuhara A, et al. Antagonistic and agonistic effects of an extracellular fragment of nectin on formation of E-cadherin-based cell-cell adhesion. Genes to Cells. John Wiley & Sons, Ltd; 2003;8:51–63.

61. Grutzendler J, Kasthuri N, Gan WB. Long-term dendritic spine stability in the adult cortex. Nature. Nature Publishing Group; 2002;420:812–6.

62. Zuo Y, Lin A, Chang P, Gan W-B. Development of long-term dendritic spine stability in diverse regions of cerebral cortex. Neuron. Elsevier; 2005;46:181–9.

63. de Vivo L, Faraguna U, Nelson AB, Pfister-Genskow M, Klapperich ME, Tononi G, et al. Developmental Patterns of Sleep Slow Wave Activity and Synaptic Density in Adolescent Mice. Sleep. Oxford University Press (OUP); 2014;37:689–700.

64. Cruz-Martín A, Crespo M, Portera-Cailliau C. Delayed stabilization of dendritic spines in fragile X mice. J Neurosci. Society for Neuroscience; 2010;30:7793–803.

65. Sheppard D. Dominant negative mutants: tools for the study of protein function in vitro and in vivo. Am. J. Respir. Cell Mol. Biol. 1994. p. 1–6.

66. Keck T, Keller GB, Jacobsen RI, Eysel UT, Bonhoeffer T, Hübener M. Synaptic scaling and homeostatic plasticity in the mouse visual cortex in vivo. Neuron. Cell Press; 2013;80:327–34.

67. Bailey CH, Chen M, Keller F, Kandel ER. Serotonin-mediated endocytosis of apCAM: an early step of learning-related synaptic growth in Aplysia. Science. 1992;256:645–9.

68. Leshchyns’ka I, Sytnyk V. Reciprocal Interactions between Cell Adhesion Molecules of the Immunoglobulin Superfamily and the Cytoskeleton in Neurons. Front cell Dev Biol. Frontiers Media SA; 2016;4:9.

69. Zuo Y, Yang G, Kwon E, Gan WB. Long-term sensory deprivation prevents dendritic spine loss in primary somatosensory cortex. Nature. Nature Publishing Group; 2005;436:261–5.

70. Xie Z, Huganir RL, Penzes P. Activity-Dependent Dendritic Spine Structural Plasticity Is Regulated by Small GTPase Rap1 and Its Target AF-6. Neuron. Cell Press; 2005;48:605–18.

71. Srivastava DP, Copits BA, Xie Z, Huda R, Jones KA, Mukherji S, et al. Afadin is required for maintenance of dendritic structure and excitatory tone. J Biol Chem. American Society for Biochemistry and Molecular Biology; 2012;287:35964–74.

72. Arikkath J, Reichardt LF. Cadherins and catenins at synapses: roles in synaptogenesis and synaptic plasticity. Trends Neurosci. NIH Public Access; 2008;31:487–94.

73. Schuman EM, Murase S. Cadherins and synaptic plasticity: activity-dependent cyclin-dependent kinase 5 regulation of synaptic beta-catenin-cadherin interactions. Philos Trans R Soc Lond B Biol Sci. The Royal Society; 2003;358:749–56.

74. Arikkath J, Peng IF, Ng YG, Israely I, Liu X, Ullian EM, et al. δ-Catenin Regulates Spine and Synapse Morphogenesis and Function in Hippocampal Neurons during Development. J Neurosci. Society for Neuroscience; 2009;29:5435–42.

75. Abe K, Chisaka O, Van Roy F, Takeichi M. Stability of dendritic spines and synaptic contacts is controlled by αN-catenin. Nat Neurosci. Nature Publishing Group; 2004;7:357–63.

76. Welsh CA, Stephany CÉ, Sapp RW, Stevens B. Ocular dominance plasticity in binocular primary visual cortex does not require C1q. J Neurosci. Society for Neuroscience; 2020;40:769–83.

