## Supplementary figures and images for "Precise levels of Nectin-3 and an interaction with Afadin are required for proper synapse formation in postnatal visual cortex"

### Additional File 2

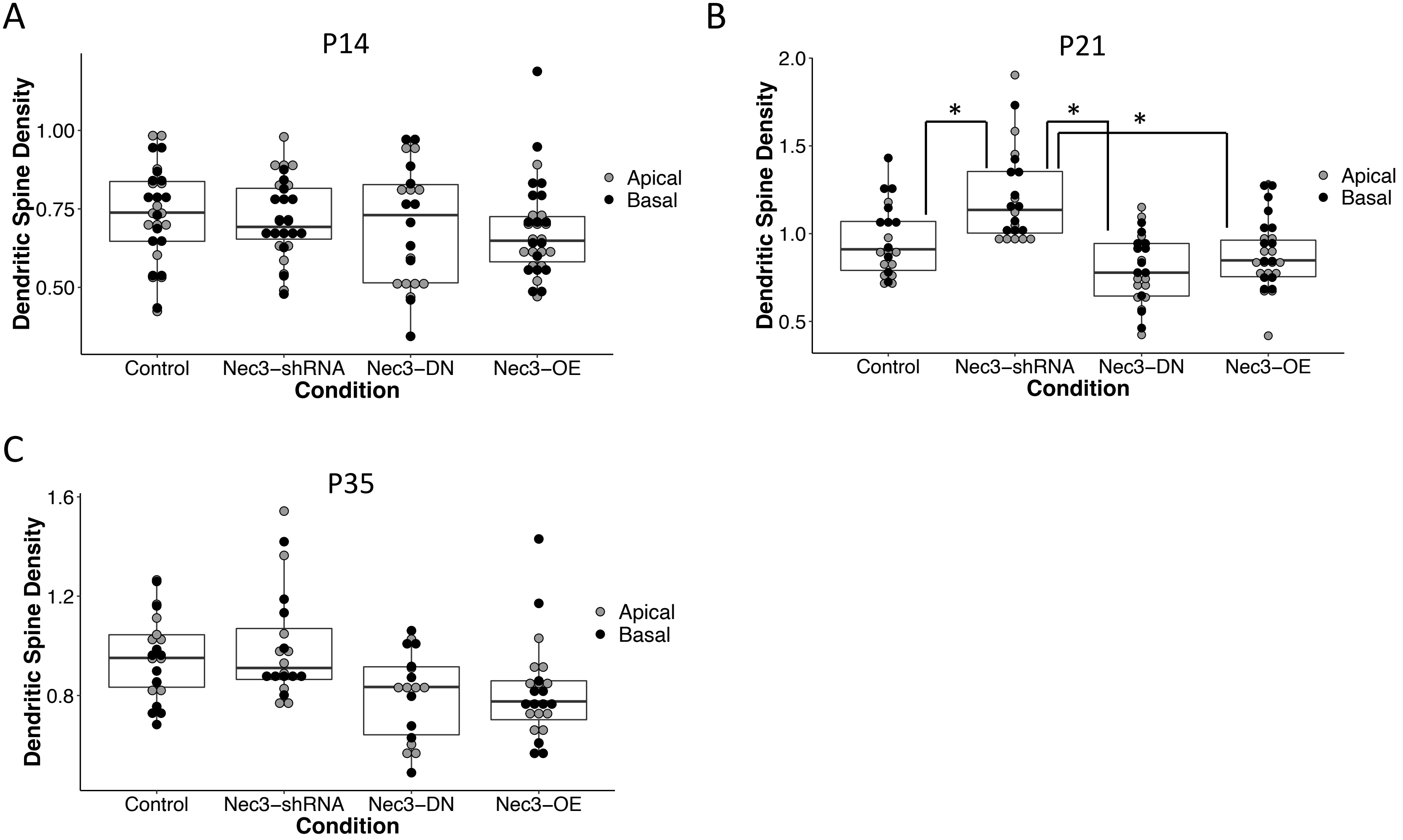
